# Public science data suggest that fish is the most important factor in reducing small pond biodiversity in Czechia

**DOI:** 10.1101/2021.03.05.434043

**Authors:** Jiri Hulcr, Demian F. Gomez

**Affiliations:** School of Forest Resources, University of Florida, Gainesville, FL, USA; Institute of Microbiology AVČR, Praha, Czech Republic; Faculty of Forestry and Wood Sciences, Czech University of Life Sciences, Praha, Czech Republic; Texas A&M Forest Service, Austin, TX, USA

## Abstract

This project tested a public-science approach to the assessment of freshwater habitat quality via simple invertebrate sampling. We combined a mobile phone application and simple instruction to children to sample 50 ponds in Central Czechia, and we analyzed the data using a standard ecological statistics approach. Despite the limitation in scope and taxonomic precision, our data revealed the same patterns as academic studies of the same topic. Specifically, we conclude that the main cause of invertebrate community decline is fish overstocking, while diverse invertebrate communities require aquatic macrophytes. Pollution detectable by children has an effect on invertebrate community structure, but a different effect than fish has, and not as statistically robust. Importantly, almost all large ponds were found overstocked with fish; therefore, they support not more diversity than small ponds, but less, and serve as ecological traps. Our findings suggest that pond conservation should focus primarily on the restoration of aquatic vegetation, and that the most effective approach will be the removal of excessive fish.

## Introduction

Central Europe boosts thousands of small, man-made water bodies scattered across the landscape and municipalities. These multi-use water sources for fish, cattle, irrigation, the local fire department, stormwater catchment, or swimming, accumulated over centuries. The result is an overlooked biological gem of Europe - a network of biodiversity hotspots in landscapes that are otherwise being subject to climate drying and overuse. Consequently, small water bodies and the organisms within them serve as a ubiquitous entryway for the public, and especially children, to learn and appreciate biodiversity in cultivated landscapes.

Unfortunately, small ponds are also an epitome of an ecosystem that has undergone rapid degradation in the recent years. Many of us can contrast our early memories of ponds teeming with life and clean water, with the reality today. Most ponds are heavily eutrophic, as indicated by green to brown color, increase in turbidity, near complete disappearance of submerged vegetation, and the absence of life (Patten et al., 1994).

The two main sources of this degradation are known. First, water pollution by chemicals and nutrients leading to eutrophication is common in Central Europe, in both large and small water bodies. Second, increased turbidity and the disappearance of plants, invertebrates and amphibians is often due the increase in fish presence. The fish species most commonly implicated with negative impact on native aquatic diversity in Central Europe is the common carp (Kloskowski, 2011) introduced to watersheds west of Danube in the Medieval ages. Additional and sometimes catastrophic consequences are attributed to newly introduced herbivorous fish (Pipalova, 2006) and to fish abundance that greatly exceeds the ecological carrying capacity of the water bodies (Zambrano et al., 2001).

Small ponds are a prime target for both biodiversity maintenance, as well as education and outreach. Ponds restoration is practical even by passive conservation, without major investments. The cessation of harmful behaviors such as pollution or fish stocking is often sufficient to reverse the degradation and allow many of the ponds to return naturally to their former status of biodiversity mini-hotspots (Peretyatko et al., 2009). The self-cleaning ability of the aquatic ecosystem is well documented, and many freshwater organisms can rapidly repopulate a water body that becomes habitable.

Pond restoration is also beneficial from the policy perspective: it is cost-effective, and readily quantifiable. One such simple quantification mechanism is tested in this paper: the use of simple biodiversity index.

### Project objectives

Our first objective was to test a method by which the public can deliver robust data on aquatic biodiversity.

Community or public participatory science, commonly also known as citizen science, has now become a mainstream in connecting the research community with the public, while delivering data which are often robust enough for research analyses and publications (Silvertown, 2009). Invertebrates are frequently the target of community science, particularly projects targeting children or youth (Ries and Oberhauser, 2015;Vitone et al., 2016). An example closely related to our project is the Big Pond Dip in the United Kingdom, a comprehensive project that involved the design of a novel biodiversity index. In its first year in 2009, the project received data from 250 participants (Rose et al., 2016).

The downside of citizen-generated data is often the uncertainty about their accuracy and repeatability. To alleviate that issue, many contemporary public science projects use electronic applications, such as online forms and databases, that exert certain degree of control over the data entry.

Our goal was to test whether using a simple, illustrated form distributed over smart phones is sufficient to allow elementary school-aged children to enter data on aquatic invertebrate diversity, and whether the data will be sufficient for serious ecological statistical analysis.

The second broader objective was to use the data to test hypotheses about causes of pond ecosystem degradation. While experts are in agreement about the two main sources of small water body degradation – pollution and fish stocking – we do not know the relative importance of the two factors. Different ponds, or different municipalities, may see the same or different main pollutants. Similarly, different aquatic organisms may tolerate pollution versus fish presence, and the two factors may lead to distinct communities.

Therefore, within the second objective we tested the degree of influence of several factors – fish presence, pollution, pond type, and vegetation – on aquatic invertebrate community composition.

## Methods

In the summer and Fall of 2020, the process described here was implemented by the senior author and a group of volunteers, mostly children and occasionally their parents, in the surroundings of Jílové u Prahy, Czechia.

Data were collected in a participatory science fashion, based on a smartphone application. We developed a data entry interface for mobile phones via NestForms^®^ (https://www.nestforms.com/) to facilitate the participation of children, and at the same time to apply quality control to the data. Figure 1 shows two representative pages of the application. There were five in total; all entry fields are listed in Table 1. NestForms^®^ was selected for two main reasons – the platform offers a simple, one-button recording of GPS location, and it allows us to illustrate individual variables for better comprehension by children. Each organismal group needed to be illustrated to allow data entry by children not aware of the scientific or even common names (Fig. 1). The data are automatically uploaded from the users’ phones to our administrator’s user account on the NestForms^®^ server from which they can be downloaded at any time and further analyzed.

**Figure 1:**
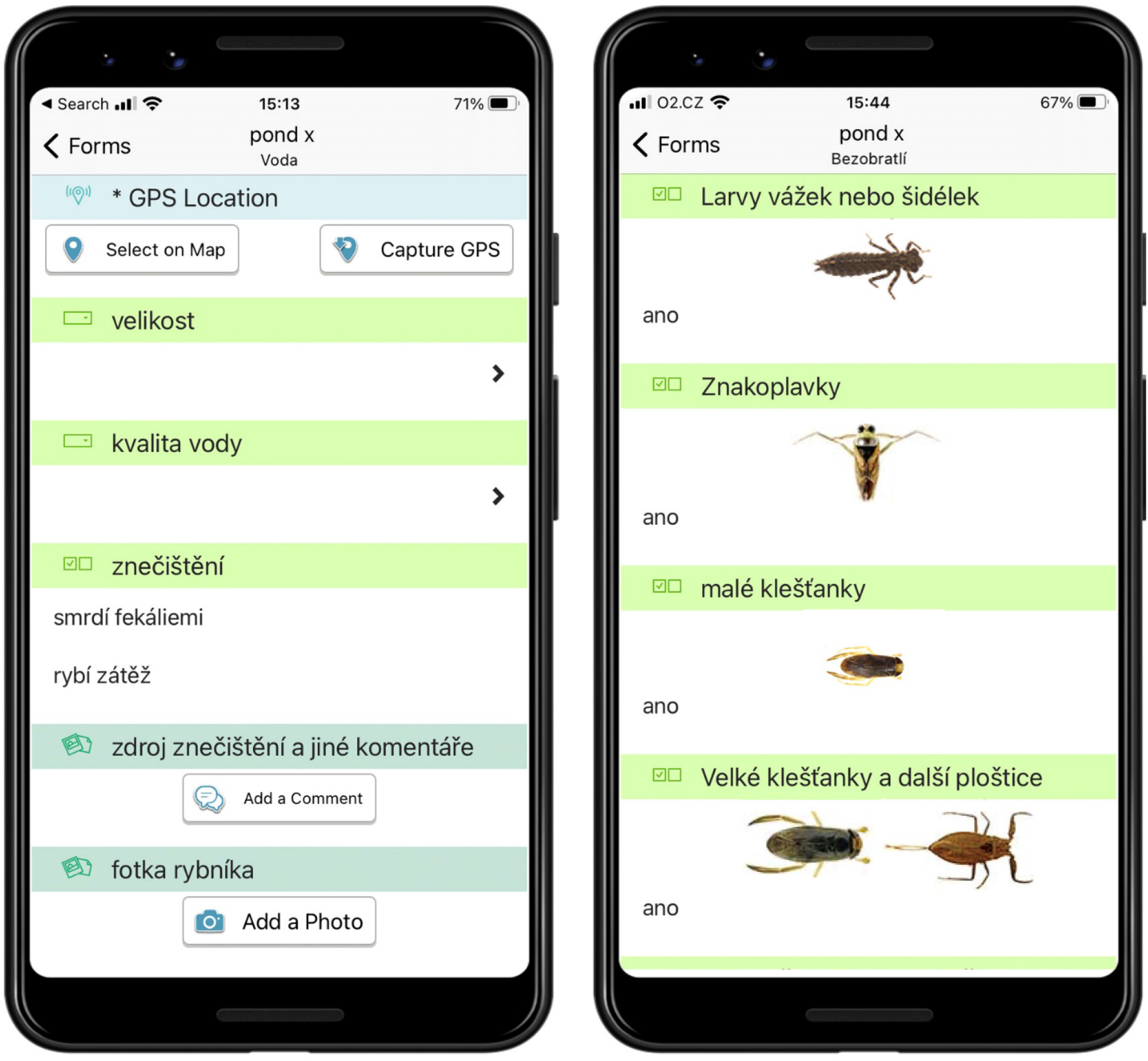
The data entry interface on a mobile phone (two out of five pages). The terms correspond to Table 1 and are written in Czech language.

**Table 1:**
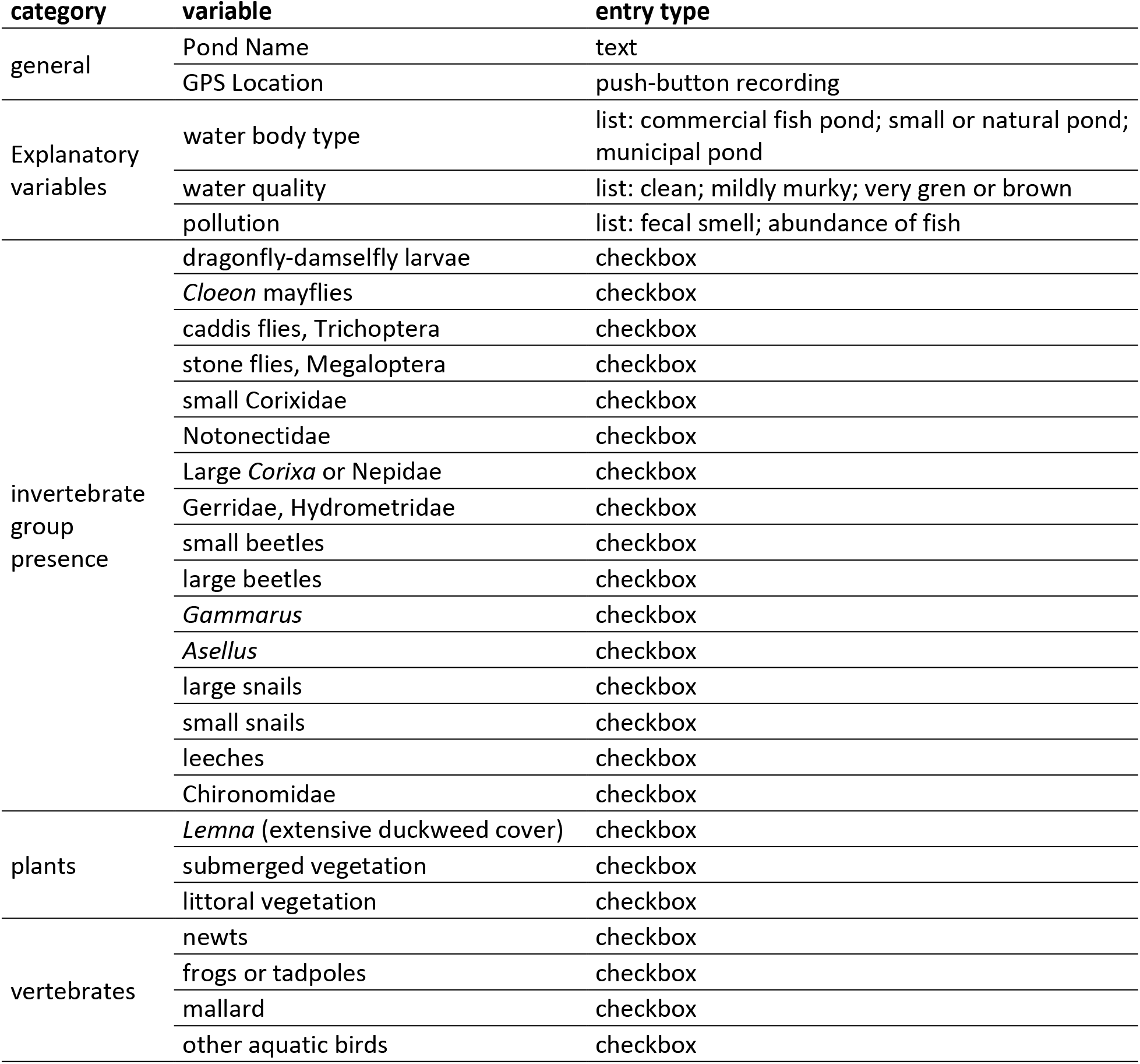
The structure of the data entry interface, implemented in the smartphone-ready form.

Entries are listed in Table 1. The environmental variables were designed to be as self-explanatory as possible, but a degree of subjectivity was inevitable in their scoring. For example, water quality was assessed simply by water color; this variable was found to be too subjective and was excluded from the subsequent analyses. The pollution source included two choices: “fecal smell present” and “abundance of fish”. The first excludes pollution without a detectable smell and therefore users may have included false negatives. Unfortunately, many ponds in municipal areas produce notable odors and their scoring as polluted was unquestionable. The choice “abundance of fish” was selected when signs of the presence of large fish were obvious – feces, artificial feeding, or the fish itself.

Our goal was to define water quality by the invertebrates captured from the ponds. This goal was similar to that of the Big Pond Dip project, with one important difference: instead of relying on an a-priori index of environmental value for each organismal group, we intended to infer the water quality directly from the data by a multivariate analysis. That is also why we opted to use a more refined list of organisms. The list of reported taxa (Table 3) is not taxonomic, but instead, it was designed with two needs in mind: ease of recognition by amateurs, and their indicator value. For example, even though water bugs (Heteroptera) could potentially be lumped in a single category, small Corixidae and the large *Corixa* were separated, since the two groups are easily recognizable and each indicates different environmental qualities. An opposite example includes small aquatic beetles which are potentially excellent as habitat quality indicators, but are impossible to recognized for a non-specialist, and therefore, they were kept in a single category.

### Date entry application

### Field collecting

The collecting took place in the vicinity of Jílové u Prahy. Users searched for water bodies on www.mapy.cz and visited each of them until 50 ponds were reached. Only publicly accessible water bodies were surveyed. Nets designed for aquatic macroinvertebrates manufactured by “Petr Dobeš– Sítě” in Jílové u Prahy (www.dobes-site.cz) were used for the sampling.

Sampling was conducted following three rules, communicated to the participants:

1. Sample every water body marked as such on the map, no matter how small or large, natural or artificial, clean or polluted. The goal was to attain a representative sample of pond qualities, not just to focus on natural ponds to maximize the collected biodiversity;
2. focus on areas with vegetation, if present, or along the sides, if no vegetation is present,
3. sample ten different spots within in each water body by a few sweeps.

### Statistical analysis

All statistical analyses were conducted in R (R_Development_Core_Team 2020). First, we tested which variables predict the greatest or the lowest species diversity. To test how the explanatory variables predict the species composition of the aquatic invertebrates, we modeled invertebrate diversity using a generalized linear model (GLM), implemented by the glm() function. The explanatory variables used were fecal smell, abundance of fish, presence of submerged vegetation, presence of littoral vegetation, and pond category. Model selection (to eliminate variables that do not explain significant amount of variation in the data) was carried out using the AIC criterion, implanted by the dredge() function of the MuMIn package.

The second goal of the analysis was to infer how individual invertebrate species correspond to environmental degradation. This we analyzed using a Redundancy analysis (RDA), implemented by the rda() function in the vegan package. The eigenvalue for each species inferred from statistically significant axes was inferred to be their indicator value. Significance of individual canonical axes was tested using a forward permutation test within the RDA analysis, with 999 permutations.

Lastly, we assessed which water body category supports the greatest or the lowest biodiversity. Water bodies were categorized as “small or natural pond”, “municipal pond” (these are typically within human inhabitation, often paved) and “large pond”. Many of the latter ones are the typical fish ponds common in the central European landscape. We modeled invertebrate diversity for pond category using a GLM with a negative binomial distribution and a log-link function. Model distribution was selected based on residual analysis. Model parameters were estimated by the glm.nb function of the MASS package (Ripley et al., 2013).

## Results

### Public engagement

Children were able to carry out the sampling and date entry after a brief initial training, but they were not able to start the data collection without adult assistance. The main reasons for this included their lack of understanding of proper net handling, lack of knowledge of the organisms, and lack of practice in estimating the environmental variables. All three hurdles were readily overcome with basic training. Becoming proficient in net handling only required two simple instructions: “sweep primarily in vegetated areas, if available”, and “sweep as hard as you can”. The practice of sweeping only ten times per pond is part of the guidelines and was understood without any training.

### Pond quality

All but three ponds yielded at least some aquatic invertebrates. The ponds which produced none were excluded from further analyses.

The model indicated that submerged vegetation, littoral vegetation and abundant fish explain significant proportion of the variation (60%). The model with the lowest AIC value of 221.38 (Table 2 and 3) retained variables submerged vegetation and littoral vegetation as positively correlated with species richness, and the variable abundant fish as negatively correlated. Fecal smell was dropped by the model selection as a variable that does not explain sufficient amount of variation.

**Table 2:** The model selected with the lowest AIC value.

| Species_Richness ~ abundant_fish + submerged_vegetation + littoral vegetation |  |  |  |  |
| --- | --- | --- | --- | --- |
|  | Estimate | Std. Error | t value | Pr(>\|t\|) |
| (Intercept) | 3.326287979 | 0.552676012 | 6.018513388 | 2.72E-07 |
| abundant_fish1 | -1.894319683 | 0.613620823 | -3.087117666 | 0.003419223 |
| submerged_vegetation1 | 2.681638045 | 0.764203188 | 3.509064196 | 0.001017601 |
| littoral_vegetation1 | 2.558784676 | 0.638777057 | 4.005755446 | 0.000223815 |

**Table 3:** Species indicator value derived from their score on the first canonical axis (the only significant one) within the RDA analysis. the lower the score, the less the organism tolerates fish and the more it is associated with undisturbed aquatic vegetation.

| invertebrate group | RDA1 score |
| --- | --- |
| small beetles (Noteridae, Dytiscidae, Hydrophilidae) | -0.311002795 |
| Gerridae and Hydrometridae | -0.302000804 |
| large beetles (Dytiscidae) | -0.241921604 |
| Chironomidae | -0.204788855 |
| dragonfly and damselfly larvae | -0.199369274 |
| *Asellus* | -0.158432181 |
| small snails | -0.105609365 |
| *Gammarus* | -0.096840497 |
| Notonectidae | -0.092545245 |
| *Cloeon* mayflies | -0.087956542 |
| leeches | -0.070516474 |
| Megaloptera | -0.065014947 |
| large snails | -0.001457765 |
| large *Corixa* or Nepidae | 0.083384702 |
| small Corixidae | 0.482777569 |

### Invertebrate communities

The first RDA axis (Fig. 2) explained 67 % of the variation, second 14%. Only the first axis was statistically significant (permutation test a_1_: p<0.001, a_2_: p=0.63). The overall pattern was similar to that seen in the model selection: the variable that was most correlated with the first axis is “abundant fish”, the variable most correlated in the opposite direction is “submerged plants”. The second axis, although not statistically significant, is highly correlated with the variable “fecal smell”, and partly correlated with “littoral vegetation”. We interpret this result such that 1) fish presence indicates most of the degradation-related species composition, 2) aquatic macrophytes predict the presence of most invertebrate diversity, 3) and nutrient pollution has a measurable negative effect on the species composition but not of the same magnitude or direction as fish presence.

**Figure 2:**
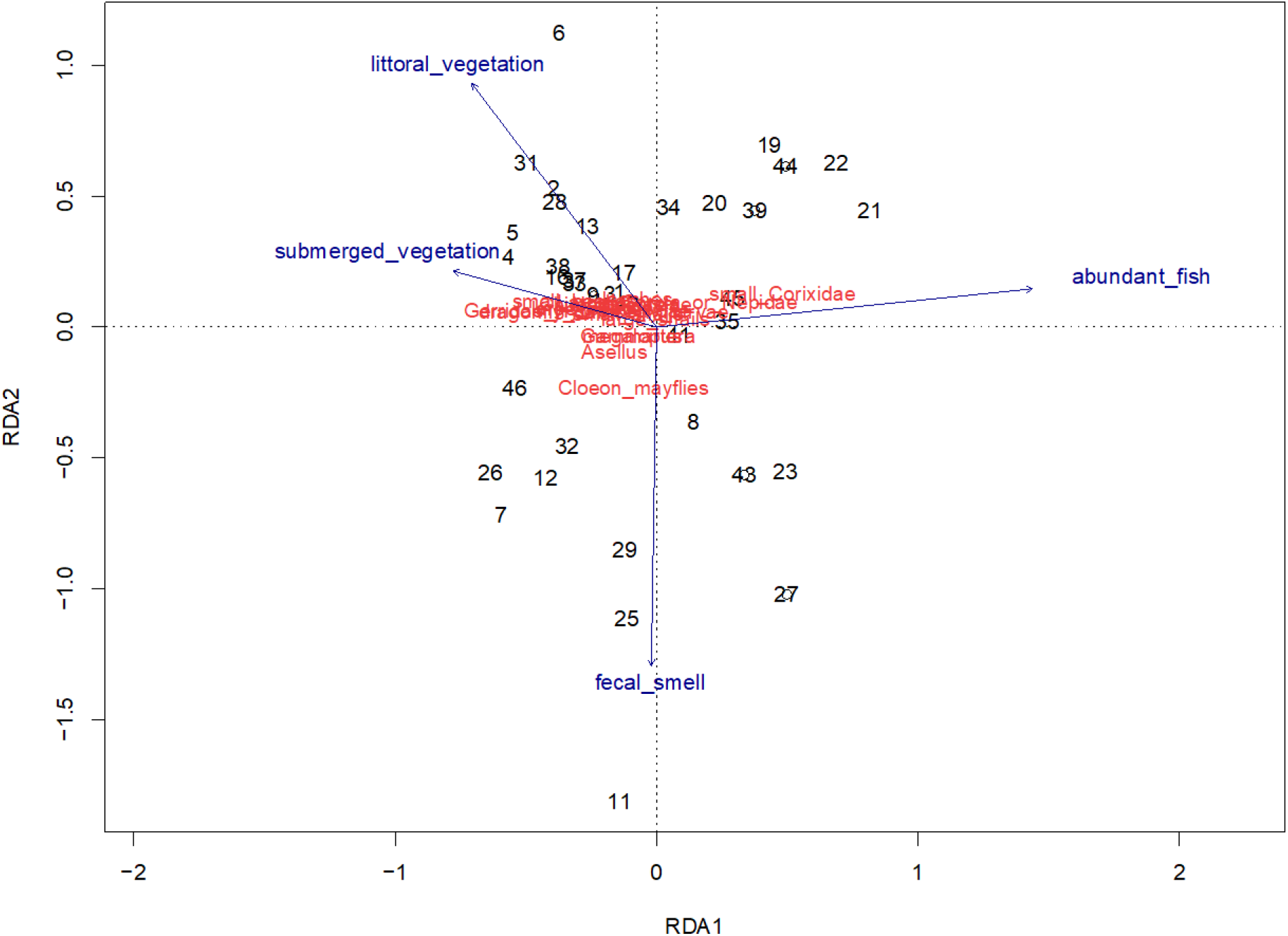
Results of the RDA analysis. Numbers refer to samples – the individual ponds – and their proximity indicates similarity in the composition of their invertebrate communities. Sample locations show compositional similarity to each other. Samples are usually dominated by species that are located near them in ordination space. Length of vectors shows the importance to the ordination. Direction indicates the correlation with each axis. Sites closer to those arrows have higher abundant fish or fecal smell, etc. Angles between axes show the correlations between environmental variables

We used the score for each species on the statistically significant first canonical axis as an their environmental indicator value (Table 3). The value essentially reflects the degree of fish load which the particular species tolerates, or the degree of plat abundance that it requires. The RDA was then also used to evaluate each pond based on the invertebrate community it hosts (Table 4).

**Table 4:**
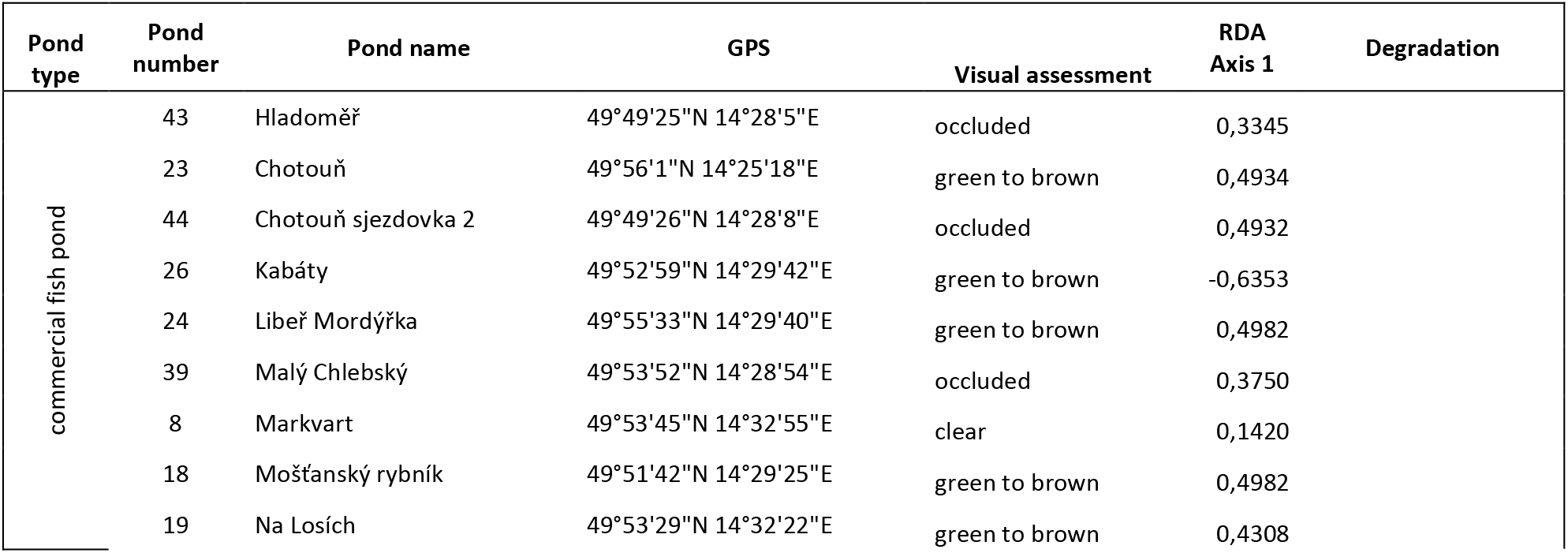
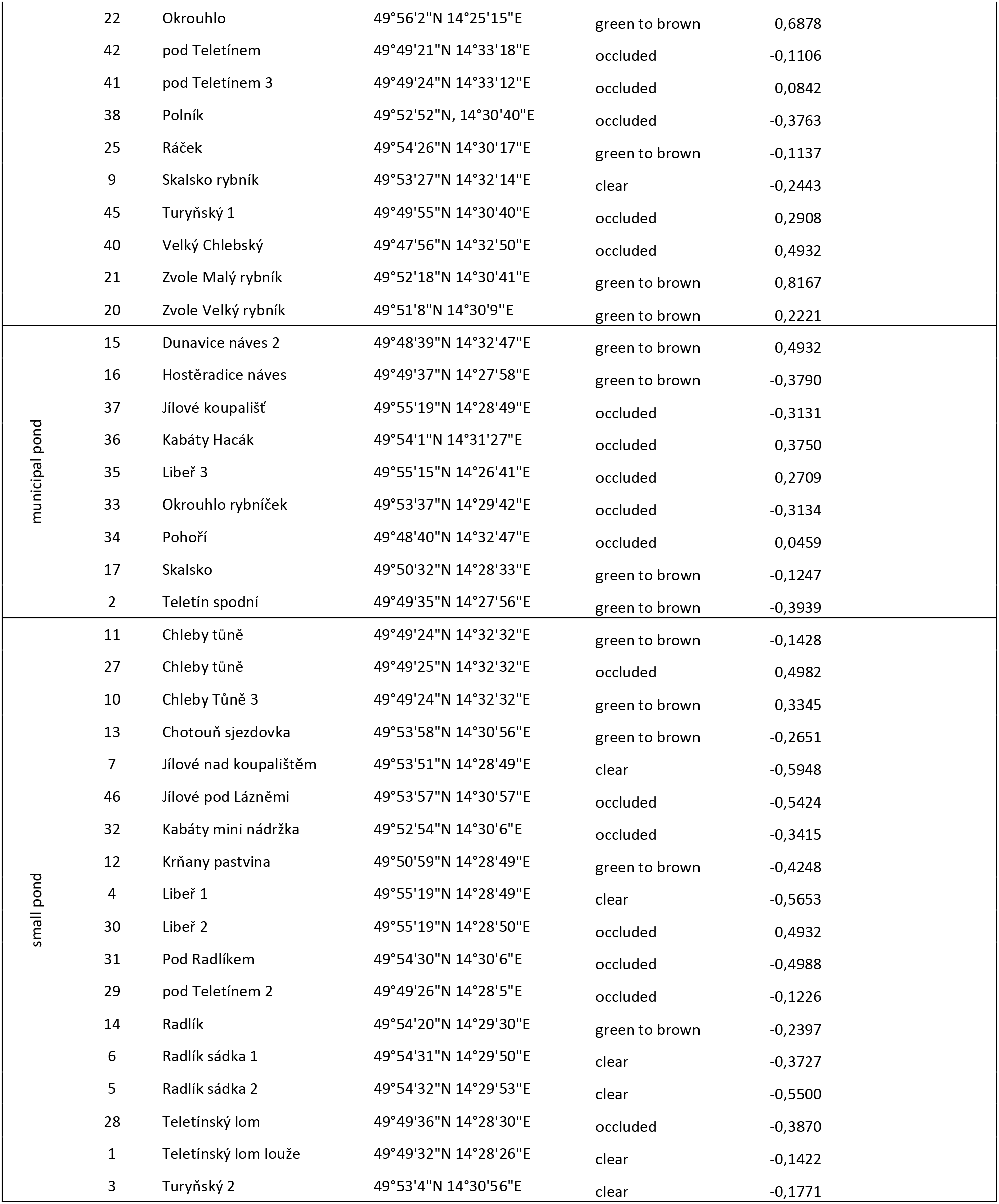
The list of ponds which yielded invertebrates, with their local names and GPS coordinates. The column Pond number corresponds to the pond IDs’ throughout the analyses and the RDA plot. Pond number reference table

The first axis, which by default explained the greatest amount of variation in the data, was most correlated with the factor “fish stocking”, and negatively correlated with “submerged vegetation”. The second axis was mostly correlated with “fecal smell” in one direction and with “littoral vegetation” in the opposite direction.

### The effect of pond type and size

Our test of effect of the pond category on the invertebrate community showed that small ponds supported a richer community than either municipal ponds or large ponds (Fig. 3), but not statistically significantly. That the richest communities reside in the category “small or natural pond” is not surprising, as nearly all the municipal ponds are paved and the large ponds were used for intensive fish production, and therefore support the most depauperate ecosystems. Typically, the only invertebrates collected in large ponds were small Corixidae.

**Figure 3:**
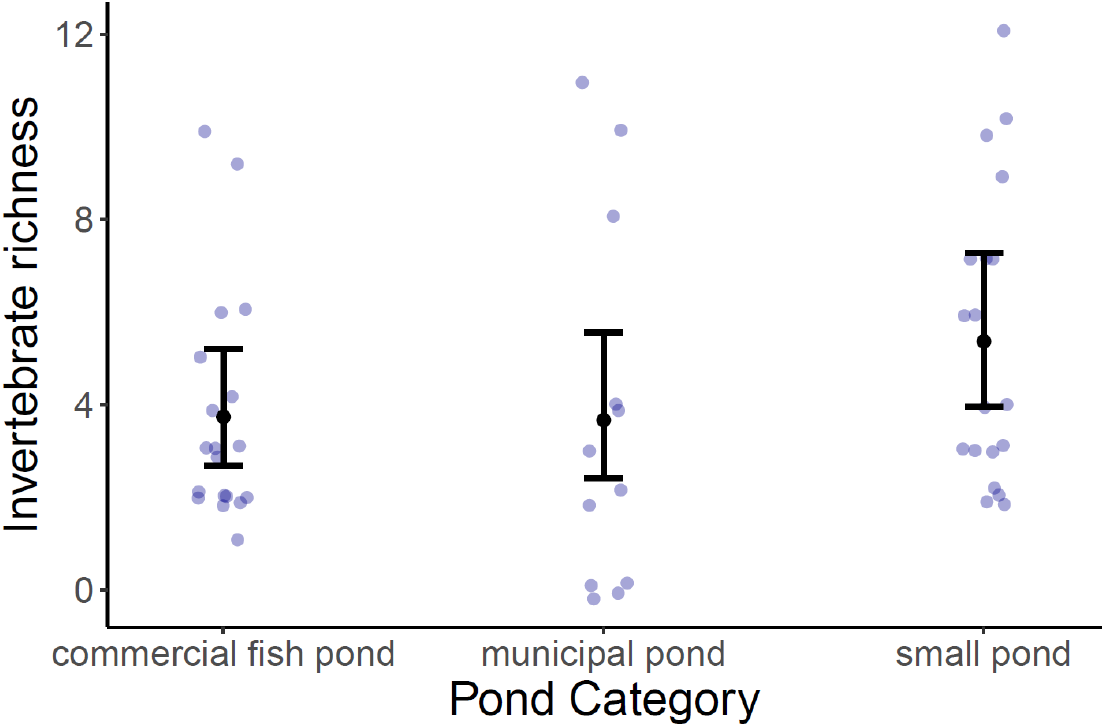
Variation in invertebrate diversity by pond size and type. Bars represent model predictions for each pond type (intervals show 95% confidence). Black circles represent the mean. Purple circles represent original observations.

## Discussion

Our results demonstrate that simple mobile electronic interface and basic training enables even youth and amateurs to perform a biodiversity survey and deliver high quality data. However, in-person introduction and mentoring was a condition of success. It is unlikely that such data would be obtainable if participants were only given written or otherwise non-interactive instructions, without the opportunity to receive feedback on, for example, assessment of the pond quality or fish presence.

Our list of organismal groups and their indicator value largely reflect those of another citizen-powered project, the Big Pond Dip. However, the indicator values attributed to some organismal groups differ in interesting ways. For example, groups such as Chironomidae and damselflies are relatively low on our scale of degradation tolerance, while they were assigned high tolerance on the Big Pond Dip scale. The reason is probably the fact that our community composition mostly reflected fish overstocking, which the British dataset may have reflected other sources of degradation, such as nutrient loading, which was not captured in our sampling.

Several organismal groups stand out at somewhat counterintuitive in their indicator value. Small Corixidae, the “water boatmen” (but not the larger species in the genus *Corixa*), were strongly positively represented in the ponds with the greatest degradation score, typically in commercial fish ponds. Similarly, the mayfly genus *Cloeon* was a surprise – despite the fact that mayflies *sensu lato* are known as indicators of pristine unpolluted aquatic ecosystem, *Cloeon* is an exception, and thrives even in severely disturbed habitats. Large snails appeared to be more tolerant of degradation than small snails in our dataset, but that was likely because of the ease of detection of the large ones, which increased their reporting probability.

The variable “fecal smell” showed an influence on the species composition that was distinct from other variables, but it was not statistically significant. This lack of strength of signal is probably due to the low rigor of our data, rather than being truly interpretable as biological fact, because water quality has been shown many times to impact species composition and diversity (Abel, 1996). Still, there are at least two interesting conclusion that can be made from our result. First, fecal smell, as an obvious sign of major aquatic habitat degradation, is not necessarily correlated with fish loading. In other words, pollution of small ponds, usually municipal or agricultural, and overstocking in commercial ponds, are two different types of environmental degradation, and they create different aquatic communities. Second, many invertebrates are able to persist even in heavily polluted and/or eutrophic water, particularly those that breathe atmospheric oxygen and do not rely on gills (Gaufin, 1973). But even these pollution-tolerant organisms disappear from waters overstocked with fish.

From the nature conservation perspective, the most important finding is that large ponds host less biodiversity than small ponds. This appears to counter one of the basic ecological rules, the species-area relationship, which posits that larger areas support greater diversity of species regardless of the ecosystem. The fact that we are seeing a reversed pattern in our pond survey is not because pond communities follow different ecological rules, but because of human intervention. Nearly all larger ponds in our focal area have been turned to massively eutrophic fish production facilities. Nearly all the municipal ponds within towns and villages are paved, devoid of vegetation and also often stocked with fish.

## Conclusions

Our preliminary data from Central Czechia indicate that, as a result of intensive fish stocking, populations of aquatic plants and invertebrates largely disappeared from the majority of large ponds. The potential of larger water bodies to provide ecosystem services is therefore not only missed, but actually reversed: fish ponds in the Czech landscape now serve as ecological traps. It is mostly the small, commercially unusable water bodies that support the remaining biodiversity heritage.

**Table 5:**
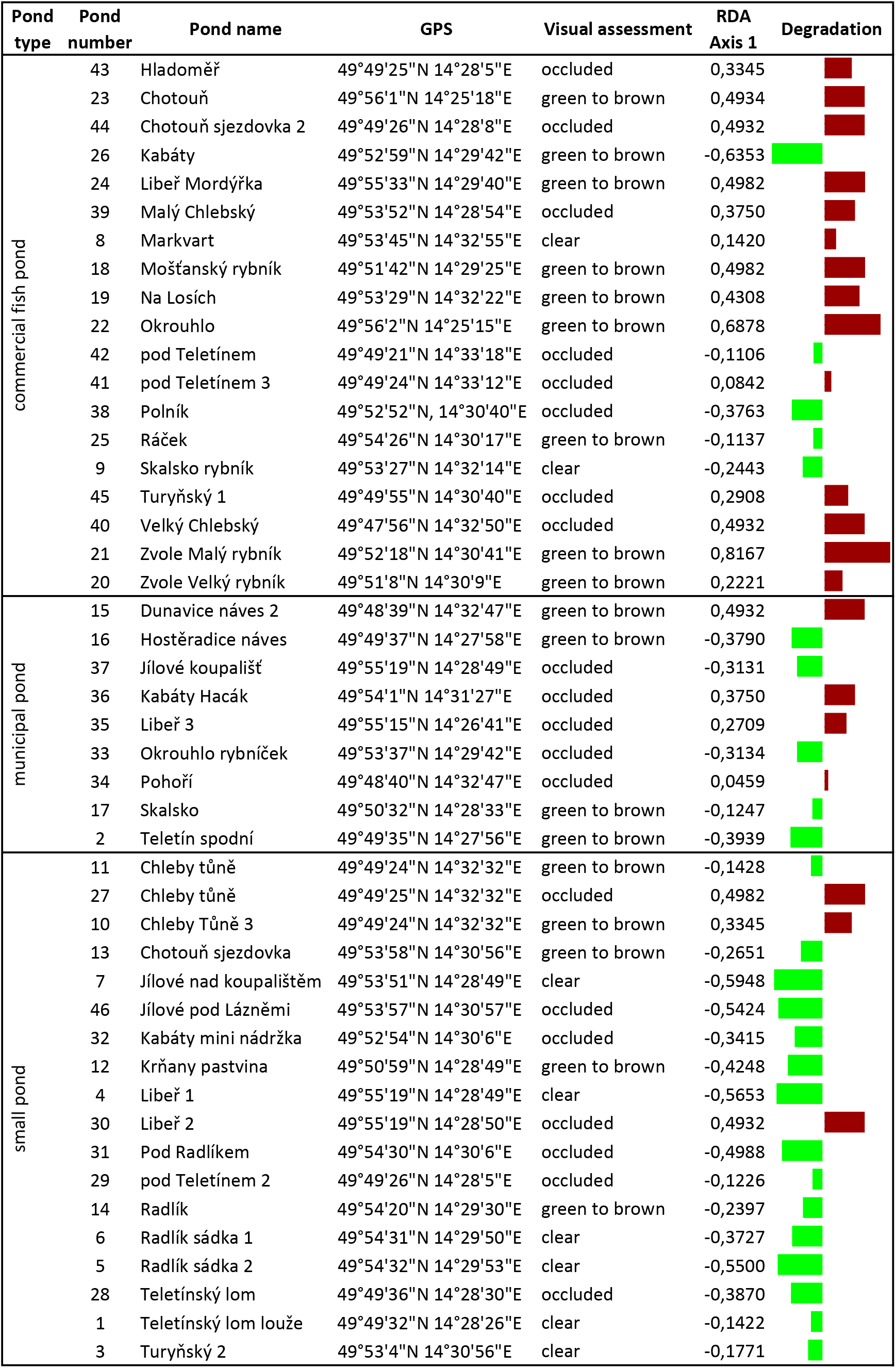
Summary of results. The “Degradation” bar displays the indicator value of each pond from the RDA (green = correlated with high invertebrate diversity, red = negatively correlated).

## Acknowledgements

We thank Sabina Lucky, Tobias Lucky, František Kolařík, Eva Kolaříková, Miroslav Kolařík, Jiří Pazour, Julie Pazourová and several other anonynmous participants for their involvement. Funding to JH was provided by the Fulbright Program of the United States, the Biological Resource Management for Sustainable Agricultural Systems of the Organisation for Economic Co-operation and Development, and the University of Florida Faculty Enhancement Opportunity fund.

